# FRONTAL AND PARIETAL CONTRIBUTIONS TO PROPRIOCEPTION AND MOTOR SKILL LEARNING

**DOI:** 10.1101/2025.10.29.685338

**Authors:** Manasi Wali, Hannah J. Block

## Abstract

Motor skill learning is the process of developing new movements with practice until they can be performed automatically. This necessitates the interaction of high-level cognitive processes with low-level sensorimotor mechanisms. Skill learning involves not only changes in the motor system but also proprioception (position sense). Proprioceptive deficits increase variability and decrease accuracy of movement. The somatosensory cortex, where low-level proprioception is processed, is known to play a role in motor skill learning, but the involvement of higher-level proprioceptive regions is unclear. Dorsolateral prefrontal cortex (DLPFC) has been linked to the high-level early stages of motor learning, and indirectly to proprioception. Supramarginal gyrus (SMG), an interface area between motor and sensory cortices, has been linked to higher-order proprioceptive processing. In this study, we asked how activity in DLPFC and SMG influences motor skill learning. Participants learned an upper limb motor skill designed to be spatially complex and dependent on proprioception: tracing a two-dimensional maze as accurately as possible within the desired speed range, using a KINARM Endpoint robotic manipulandum. Proprioceptive acuity (sensitivity and bias) was assessed before and after continuous theta burst transcranial magnetic stimulation (cTBS) was applied to inhibit activity in DLPFC, SMG, or Sham. To measure motor skill, movement accuracy and variability were examined at the trained speed as well as at a faster (more difficult) and a slower (easier) speed. Skill was assessed before and after cTBS, after 40 training trials (early learning) and after another 80 trials (late learning). All three groups showed improvements in movement accuracy and variance, indicating they learned the maze tracing skill. However, the Sham group improved movement variability at the faster speed significantly more than the DLPFC or SMG groups did. This suggests that DLPFC and SMG are important for the more challenging aspects of motor skill learning, consistent with their association with higher-level proprioceptive function.

## INTRODUCTION

Motor learning is the process of developing new movements with practice until they can be performed automatically (Kitago & Krakauer, 2013; Schmidt & Wrisberg, 2008). It is generally divided into altering movements in response to perturbations (motor adaptation) (Caithness et al., 2004; Krakauer et al., 1999, 2000) or acquiring new movement patterns without the presence of perturbations (skill learning) (McGrath & Kantak, 2016; Shmuelof et al., 2012). Although there is no set definition of what qualifies as a complex skill in the upper limb, our group and others have suggested that it entails synchronizing and sequencing multiple arm movements with spatial and temporal constraints (Kantak et al., 2018, 2017; Mirdamadi & Block, 2020, 2021). Practice makes the movement kinematics smoother, which indicates improvement in quality of movement and better planning (Kantak et al., 2018; Shmuelof et al., 2012). Motor skill learning is characterized by two stages: Fast and slow. The fast stage occurs when large improvements in performance occur in a relatively short period of time. The slow stage occurs later, and the improvements can be seen in the form of further gains which are achieved more slowly (Dayan & Cohen, 2011). Motor skill improvement is characterized by a change in the speed-accuracy function, meaning execution of motor patterns occurring more accurately at one or more different speed ranges (Kantak et al., 2017; Reis et al., 2009). A review by Dagmar Sternad (2018) drew attention to variability and noise during motor performance being informative biological features in understanding movement control. Skill improvement is also manifested by individuals becoming less variable over time without sacrificing speed or moving faster without an increase in variability. The authors mention that mastering a new skill involves increasing accuracy and decreasing variability or “with maximum certainty and a minimum outlay of time or energy” (Sternad, 2018). A complex neuroanatomical architecture supports the learning and retention of motor skills. Learning motor skills necessitates the interaction of several high-level cognitive processes with low-level sensorimotor mechanisms (Krakauer & Mazzoni, 2011). Different cortical and subcortical brain activations take place during the two stages of motor learning. The fast stage, i.e. the early stage of learning, is associated with activity in the frontoparietal regions such as dorsolateral prefrontal cortex (DLPFC), primary motor cortex (M1), pre-supplementary motor area (pre-SMA), the premotor cortex, supplementary motor area (SMA), parietal regions, cerebellum and striatum (Dayan & Cohen, 2011; Floyer-Lea & Matthews, 2004; Halsband & Lange, 2006). In contrast, during the slow or later stage of learning, there is an increase in the activation in M1, primary somatosensory cortex, SMA, and subcortical regions like cerebellum and basal ganglia (Floyer-Lea & Matthews, 2005; Halsband & Lange, 2006). There is also a decrease in the activation in lobule VI of cerebellum (Lehéricy et al., 2005). As skill learning progresses from early to late (fast to slow) stages, activation shifts from anterior regions to more posterior regions (Floyer-Lea and Matthews, 2005).

Motor skill learning involves not only changes in the motor system but also depends on sensory systems. One of the sensory modalities critically important for motor skill is position sense or proprioception—the ability of an individual to integrate sensory inputs from proprioceptors in muscles and joints to determine the positions and movements of body segments in space (Han et al., 2016). Proprioceptive information about the hand’s position, acquired from the peripheral nervous system, is taken into consideration by the motor system as feedback which is required for smooth and skilled movement execution (Bossom, 1974; Jeannerod, 1988). There is ample evidence that proprioceptive training can enhance motor learning. Wong et al. (2012) found that the group of subjects that were provided with additional proprioceptive training learned the desired movements better (Wong et al., 2012). A review by Aman et al. (2015) showed converging evidence that proprioceptive training overall results in 52% enhancement of motor function throughout every outcome measure (Aman et al., 2015). Cherpin et al. (2019) suggested that their results pointed towards an association between motor performance and proprioceptive deficits in chronic stroke survivors with proprioceptive impairments. Impairment in proprioception has also been associated with poor motor recovery post stroke (Cherpin et al., 2019; Findlater & Dukelow, 2016)

Many studies have shown that an enhancement in proprioception results in improvements in motor learning/decline in motor deficits or vice versa, although the neural basis of such interactions is unclear. Somatosensory cortex is the most common brain region studied with respect to proprioception and skill learning since it is associated with proprioceptive changes (Mirdamadi & Block, 2020). Cerebellum is another structure which had recently been found to be involved in changes in proprioception related to skill learning; Cantarero et al. have shown that excitatory tDCS on the cerebellum results in the enhancement of skill learning through online changes (Cantarero et al., 2015). Recently, Mirdamadi & Block (2021) determined that inhibiting S1 and cerebellum excitability disturbed proprioceptive function during skill learning (Mirdamadi & Block, 2021). Other neural substrates of interest in connecting proprioceptive function to motor skill learning are Dorsolateral Prefrontal Cortex (DLPFC) and Supramarginal Gyrus (SMG). Research has found an indirect connection between DLPFC and proprioception. Stagg et al. (2013) applied anodal tDCS over left DLPFC and found an enhanced functional connectivity between sensorimotor cortex and DLPFC using MRI. They suggested that the enhanced activity in the DLPFC might be linked to enhanced proprioception (Stagg et al., 2013). Another important region is posterior parietal cortex (PPC), in addition to acting as an interface between the sensory and motor cortex by integrating the multisensory signals with motor related information, it generates signals that are critical for updating body representation (Prevosto et al., 2011). A study by Ben-Shabat et al. (2015) found a key role of the right SMG in proprioception. They indicated that brain activation, which was higher-order proprioception related to somatosensory cortex, included the SMG. They found that in the presence of proprioceptive deficits, the right SMG’s activity was reduced. They also found that patients with stroke were susceptible to deficits in proprioception if their right SMG was affected particularly (Ben-Shabat et al., 2015). They explained that the decrease in right SMG’s function during proprioceptive impairment might be associated with the role that this region plays in spatial processing (Stephan et al., 2003). SMG is part of the somatosensory association cortex, playing a role in perception of space and location of limbs (Stephan et al., 2003) as well as the interpretation of tactile information (Goble et al., 2012; Naito et al., 2005). However, it is unclear how DLPFC and SMG modulate proprioceptive functioning in the context of motor skill learning.

The sensorimotor network associated with proprioception and motor skill learning is not fully understood, which limits the development of neurorehabilitation techniques and protocols. Neural resources like DLPFC and SMG have not been investigated in terms of their modulatory effects on proprioception and then in turn motor skill learning. The aim of the present study was to examine the role of DLPFC and SMG in proprioception and skill learning using repetitive transcranial magnetic stimulation (rTMS). This technique has been used to understand the interactions between brain and behavior and investigate the probable cause-effect associations between particular aspects of behavior and modified brain activity in a specific region (Cohen et al., 1997; Reis et al., 2008). A temporary reduction in cortical excitability can be achieved through continuous theta burst stimulation (cTBS), a patterned form of rTMS (Huang et al., 2005). Three groups of participants received cTBS over DLPFC, SMG or SHAM, before training on a maze - tracing motor skill with their dominant right hand. Proprioception was assessed before and after cTBS, and motor skill (movement accuracy and variability) was assessed at multiple time points at both trained and untrained speeds.

## METHODS

### Participants

#### Sample size and power

A total of 79 right-handed individuals (26-27 in each group, 37 males and 42 females) aged between 18-45 years (23.6 ± 5.26 years, mean ± SD) were enrolled. Participants were free of neurological or musculoskeletal problems or risk factors for TMS (Rossi et al., 2009) or MRI. All participants were right-handed, and their handedness scores were recorded using Edinburgh Handedness Questionnaire (Oldfield, 1971). The participants gave written informed consent, and protocols were approved by the Institutional Review Board (IRB) of Indiana University Bloomington.

### Experimental Design

The study took place in up to three sessions. In the first session (familiarization) we acquainted them with the study techniques, which allowed us to make sure they can tolerate TMS. If the participant was eligible after the familiarization, then they were randomly assigned to a group. Depending on group assignment, subjects received an anatomical MRI scan of their brain in a separate session. During the main session, participants performed the behavioral assessment tasks first, followed by the cTBS intervention as per group assignment. Next, they were asked to perform the behavioral assessment tasks again followed by skill training and additional assessments.

#### Familiarization session

Subjects were seated on a chair and asked to wear a pair of goggles which enabled us to accurately position the TMS coil on their heads relative to the template (Brainsight neuronavigation system). Electrodes were placed on their right index finger muscle first dorsal interosseus (FDI) to measure muscle activity upon stimulation (electromyography, or EMG). Single TMS pulses were applied over the motor area that represents the index finger. The area in the brain that elicited the strongest and most consistent muscle response in the EMG was identified as the motor hotspot. According to existing standards, we determined the resting motor threshold (RMT) at the motor hotspot as the lowest TMS intensity which produced a muscle twitch of 50 microvolts or above at least 10 out of 20 times (Rossini et al., 2015). We also performed 10 seconds of continuous theta burst stimulation (cTBS) on the two intervention areas (DLPFC and SMG) to check for each subject’s tolerance.

This session helped us to ensure that the subjects were comfortable with TMS and the TMS measurements were possible before calling them in for the main session. During this session, participants were also given some practice for the proprioception and motor tasks. Next, the subjects were randomly assigned to one of the groups in the experiment: Dorsolateral Prefrontal cortex (DLPFC), Supramarginal gyrus (SMG), and Sham (Inactive control). To locate subject-specific target positions for the participants in the DLPFC and SMG group, anatomical brain scans were taken of each subject’s brain. These scans were uploaded to Brainsight where we marked the anterior and the posterior commissure (AC and PC). Using these landmarks, Brainsight rotates the brain image to align it along the AC-PC plane, correcting for tilt and twist rotation. The overall size of the brain was found by moving the walls of a box to the edges of the brain in the lateral, vertical, and anterior-posterior (AP) direction. These distances were compared to the width of the reference brain to calculate the correct scaling factors in the three directions. Brainsight linearly transforms (translates, rotates and scales) the MNI brain to fit the scan. It enables a target to be defined in MNI coordinates, which were then used as the basis of TMS coil positioning for cTBS in the SMG and DLPFC groups.

#### Main Session

During the main experimental session, we first assessed the baseline proprioception and speed-accuracy function for each subject (Pre timepoint). This was followed by the cTBS intervention which was delivered using the Magstim Super Rapid Plus stimulator with a D70^2^ 70 mm figure of eight shape coil (Magstim Company LTD, UK). RMT was determined similarly to the familiarization session. cTBS, delivered at 70% of RMT, consisted of sets of three pulses at 50 Hz repeated at 5Hz for 40 seconds (Huang et al., 2005). It was delivered over either the left DLPFC (MNI: [-40 28 18]) or right SMG (MNI: [44 -50 46]) to modulate the hemisphere that pertains to the target hand experiencing the motor training (Fig 1A). For the sham group, an unplugged TMS coil was kept on the M1 target used for finding the RMT and a second coil was held, out of the participants’ view, behind their head, plugged in to make sure the sound of the cTBS was audible. cTBS was followed by another set of proprioceptive and motor skill assessments (Post 1 timepoint). Subjects next performed a short interval of motor training (40 trials), another motor skill assessment (Post 2 timepoint), and a long interval of motor training (80 trials). The session concluded with a final motor skill assessment (Post 3 timepoint) and some questions about the subject’s subjective experiences of the procedures (Fig.1B).

**Figure 1.**
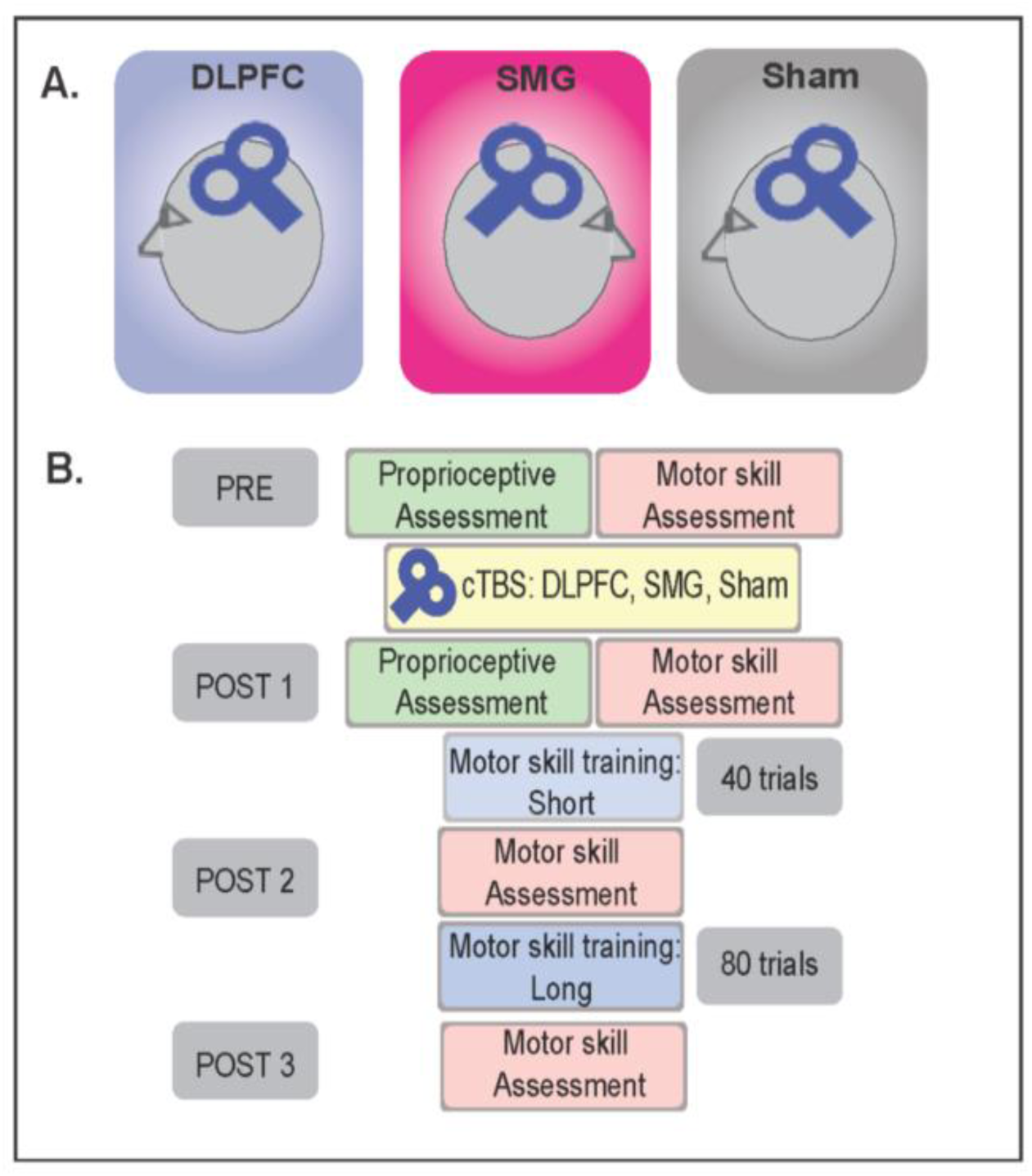
**A.** Illustration of the stimulation targets depicting the three different groups. We delivered the continuous theta burst stimulation (cTBS) over the left dorsolateral prefrontal cortex (DLPFC), right supramarginal gyrus (SMG), or sham stimulation over the right primary motor cortex. **B.** Experimental Design. Motor skill assessment and Proprioceptive function assessment were measured by Pre and Post (Post1) intervention (cTBS delivery). The behavioral assessment was followed by short motor training (40 trials), motor skill assessment (Post2), and long motor skill training (80 trials). The task ended with another motor skill measurement (Post3). Motor skill was assessed at three different speed ranges for evaluating the speed-accuracy tradeoff function and movement variability.

#### Apparatus

Participants were seated in front of a two-dimensional (2D) virtual reality apparatus for all tasks. They grasped a KINARM End-Point robotic manipulandum (BKIN) with their right hand. The task display was viewed in a mirror, such that it appeared in the plane of the manipulandum. The mirror and a bib around their shoulders prevented direct vision of their hands, arms, and the robotic handle (Fig 2A).

**Figure 2.**
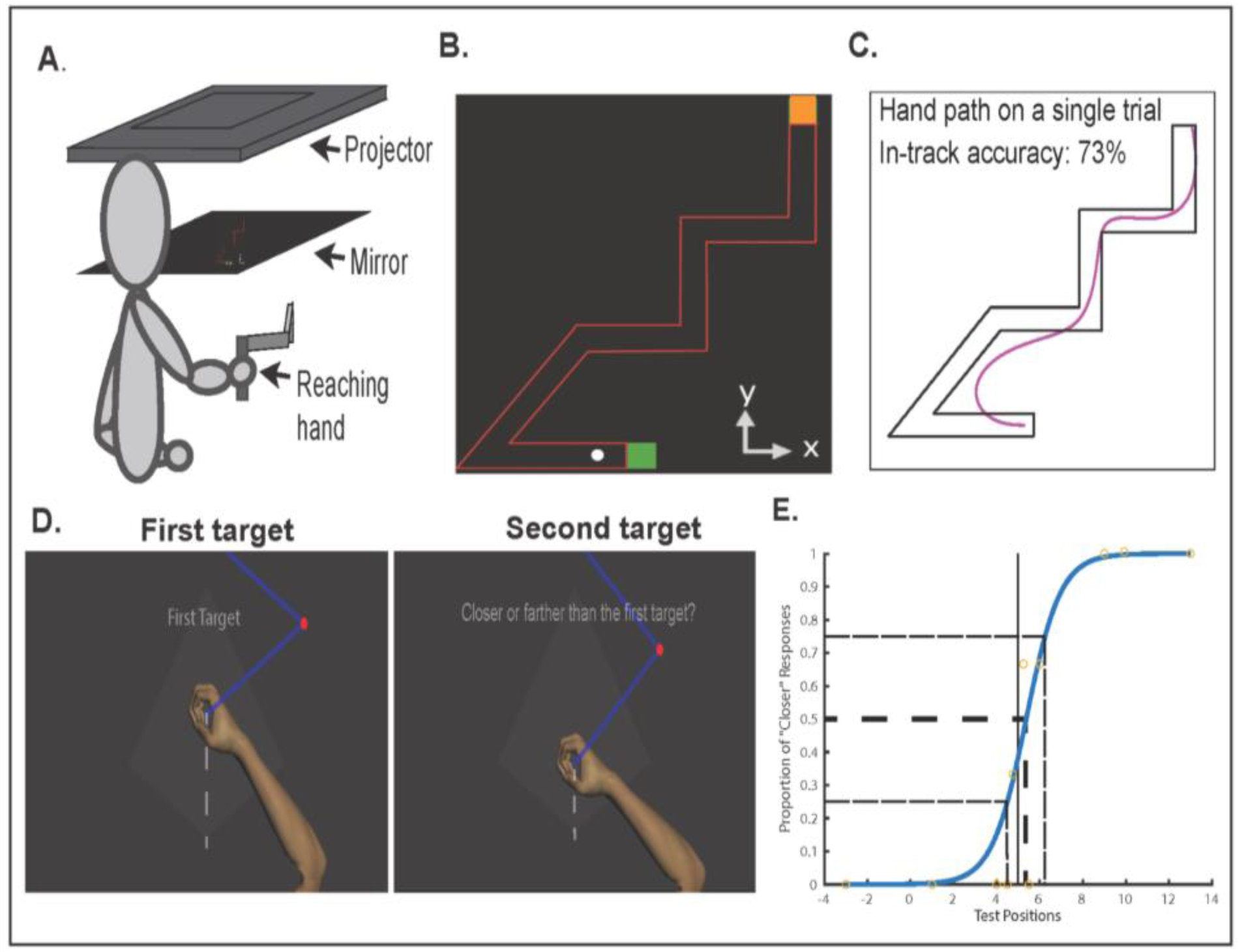
**A.** Illustration of a 2D virtual reality apparatus setup. The tasks were performed by the subjects with their right hand grasping the manipulandum handle, and vision was occluded. **B.** Depiction of the maze track used to perform the motor skill assessment task and skill training. Participants were centered with the track and seated in the negative y-axis direction. White cursor was a veridical representation of their hand movement. Subjects navigated through the maze from the start box (green) to the target (orange). **C.** Example of movement trajectory on a single trial used to compute the in-track accuracy and movement variability. **D.** Proprioceptive assessment – bird’s eye view. The participants’ hand and movement trajectory were not visible to them. The robot moved their hand to two different positions in the sagittal workspace. At the second target participants reported verbally whether their hand was closer or further to them as compared to the first target. **E.** Example proprioceptive data fitted with logistic function. The 50% point of the fitted function was defined as proprioceptive bias, and the difference between the 25% and 75% points of the fitted function was defined as the sensitivity.

### Motor skill assessment

The participants were asked to navigate through an irregular maze (1.5 cm wide) with the robotic manipulandum. The hand feedback was given veridically using a visual cursor (10 mm wide, white circle). This design was similar to Mirdamadi & Block, 2020 with six straight line segments connected abruptly (Fig 2B). Subjects were asked to enter an orange square at the start of the trial. The square turned green after 1 second, and the maze track appeared with another square at the end. The participants were instructed to reach the second target at the end of the track by moving the white cursor within the track as accurately as possible. Once the subjects reached the second target, the trial was complete. Feedback was given about their movement time by the target color (red: too fast, yellow: too slow, green: good speed) and accuracy with points. The maze was performed over three movement time (MT) ranges to assess the speed-accuracy tradeoff in three blocks of trials (MT1: 300-600 ms; MT2: 600 – 850 ms; MT3: 850 – 1100 ms). MT2 is a comfortable speed, while MT1 is more challenging and MT3 is easier. 10 trials were performed in each MT range, and the block order was randomized. The skill assessment took about 5 minutes.

For each of the three MT ranges, we calculated the in-track accuracy (movement trajectory percentage inside the track) and variability (RMSE) in each trial. Only trials in the correct MT range were analyzed. Mean in-track accuracy and variability were calculated separately for each MT bin in the skill assessment block.

### Motor Training

Practice of the maze task occurred at a single MT range (MT2). During the short motor training interval, participants trained for 40 trials. The long motor training interval consisted of 80 trials. Good speed trials were rewarded with 5 points, and additional points were given for higher accuracy levels. The motor training took about 5 minutes for short and 10 minutes for long training.

### Proprioception Assessment Task

Hand position sense was assessed by a constant stimuli task (Simpson, 1988) using passive movements. Participants were asked to move the handle to a starting position. Next, the robot moved their hand to the first unseen position (first target), followed by randomly moving it either up or down (distractor movement) and then to another unseen position (second target). Participants were asked to verbally report whether the second target was closer or further to them as compared to the first (reference) target. There was a total of 78 trials, which took about 12 minutes. This assessment was performed at two timepoints (Pre, Post1) (Fig 2D).

We calculated the proportion of trials that the participant responded “closer” across the different test positions and fitted the data with a logistic function. The 50% point of the fitted function (perceptual boundary) was defined as the bias or point of subjective equality (PSE) (Wood et al., 2023). This was the point at which the target equally felt closer or further. We also calculated the sensitivity or just noticeable distance (JND), which was the distance between the 25% and 75% points of the fitted function (Han et al., 2016) (Fig 2E).

### Statistical Analysis

To assess changes in motor skill, we did a mixed-model ANOVA on movement accuracy and movement variability (RMSE) for each speed range (MT1, MT2, MT3) with between-subject factor Group (DLPFC, SMG, Sham) and within-subject factor Timepoint (Pre, Post1, Post2, Post3). To control for individual differences at baseline, we repeated these analyses after subtracting the Pre timepoint from all the Post timepoints (Post1-Pre, Post2-Pre, Post3-Pre).

Changes in proprioception across the two timepoints could take the form of changes in proprioceptive sensitivity (JND) and/or proprioceptive bias (PSE). For each of these outcome measures, we did a two-way ANOVA with Timepoints as within-subject factor (Pre, Post1) and groups (DLPFC, SMG, Sham) as between subject factor, with appropriate post-hoc tests upon significant interactions (p<0.05).

To examine the performance of the three groups on motor training short and long, we did a mixed model ANOVA with within-subject factor Trial and between-subject factor Group (DLPFC, SMG. Sham). Lastly, to see whether the RMT differed across groups, we performed a one-way ANOVA on RMT with between-subject factor Group (DLPFC, SMG, Sham).

Statistical analyses were performed using JASP (version 0.18.3). For each ANOVA, we used the Shapiro-Wilk test and Kruskal-Wallis test to check for assumptions for normality and homogeneity of variance. Any significant effects were supported by post-hoc tests and corrected for multiple comparisons using Tukey’s HSD method. Alpha was defined as 0.05 for all hypothesis tests.

## RESULTS

### Motor Training

Although movement accuracy generally improved across both the short and long training intervals, there was no evidence that the three groups performed differently during training. We did a mixed model ANOVA on movement accuracy in each training interval with factor Groups (DLPFC, SMG, Sham) x Trials. In the short phase of training (Fig. 3A) there was a main effect of Trial on movement accuracy (F(39,2964) = 1.980, p<0.001), consistent with improved performance across this period. However, there was no Trial x Group interaction (F(78,2964) = 0.428, p = 1.000) or group difference (F(2,76) = 0.628, p = 0.536). Similarly, during the long phase of motor training (Fig. 3B) there was a main effect of Trial (F(79,6004) = 4.156, p < 0.001) but no Trial x Group interaction (F(158,6004) = 0.855, p = 0.905) or group effect (F(2,76) = 0.615, p = 0.543). As an additional check for group differences during motor training, we compared average accuracy in the first block of training (short training Block 1) to last block of training (long training Block 8) across groups (Fig. 3C). There was a main effect of Block (F(1,76) = 4.268, p = 0.042), consistent with improved performance across training. However, there was no interaction effect between Block x Group (F(2,76) = 0.425, p = 0.655) or group effect (F(2,76) = 0.826, p = 0.442), meaning all the groups improved similarly.

**Figure 3.**
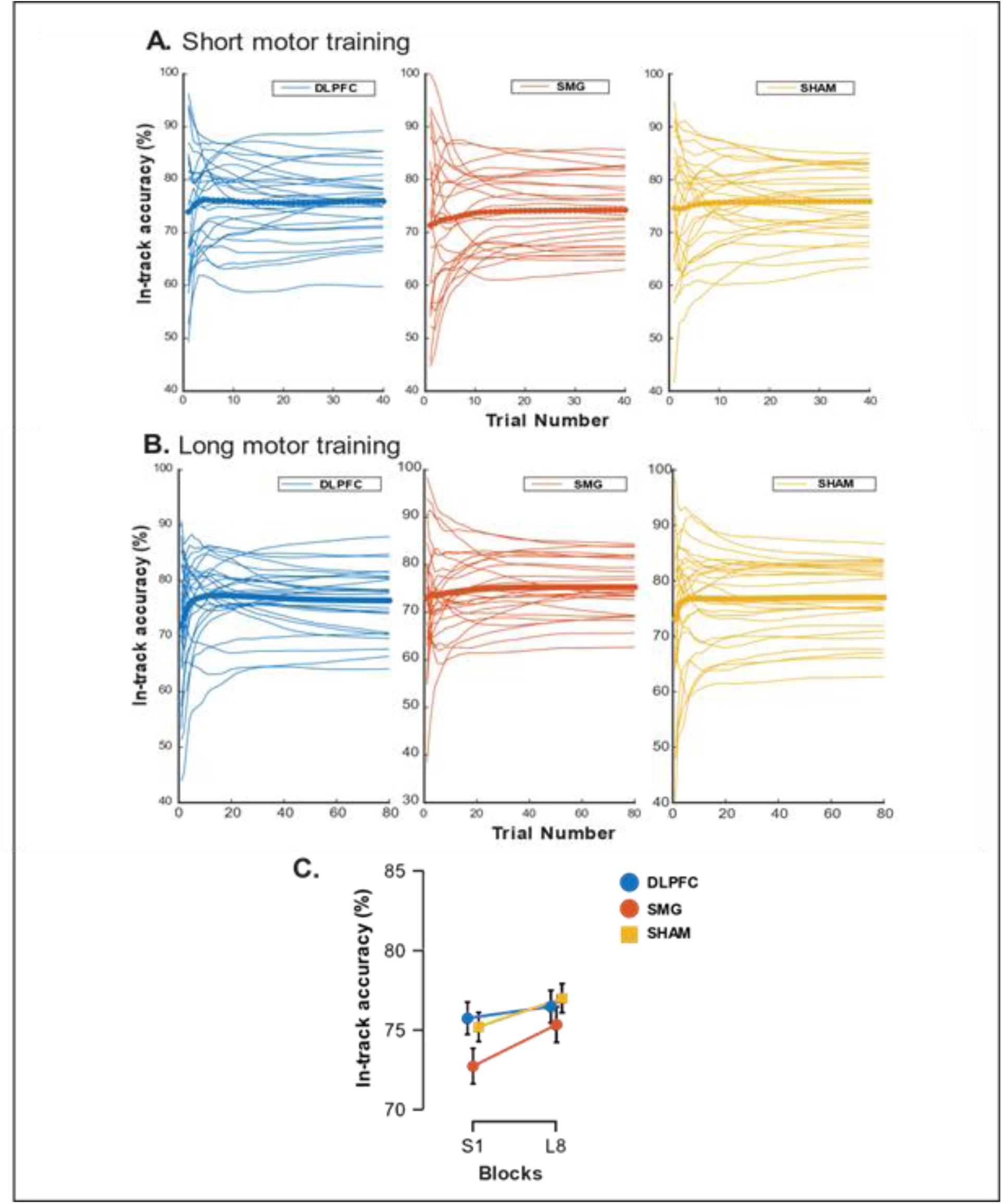
In-track accuracy percentage during short (40 trials) and long (80 trials) motor training. Thin lines indicate individual subject performance in each group. Performance seemed to plateau within the first 15 trials, with substantial variability in individual subjects’ performance. **A.** Accuracy percentage for each group (DLPFC, SMG and Sham) during short motor training at each movement trial at the medium speed range. **B.** Accuracy percentage for each group (DLPFC, SMG and Sham) during long motor training at the medium speed range. **C.** Improvement in performance from the first block in short training to the last block in long training. Each block represents an average of 10 trials.

### Motor Skill Learning

#### Movement Variability

We performed a mixed-model ANOVA on movement variability (RMSE) with three groups (DLPFC, SMG, Sham) x four timepoints (Pre, Post1, Post2, Post3) at each speed range. At MT1 we found a significant timepoint main effect (F(3,225) = 9.856, p<0.001), consistent with improved performance across time, but no Timepoint x Group interaction (F(6,225) = 1.849, p = 0.091). There was a group effect (F(2,75) = 4.417, p = 0.015), with post-hoc tests showing that the DLPFC group’s variance was less than the SMG group’s (p = 0.011) on MT1 speed range. At speed range MT2 we found a significant main effect of timepoint (F(3,228) = 5.903, p<0.001) but no Timepoint x Group interaction (F(6,228) = 0.891, p = 0.502). There was no group effect (F(2,76) = 1.324, p = 0.272). In other words, variability did not differ across groups for speed range MT2. For MT3 we found a significant timepoint main effect (F(3,225) = 3.703, p = 0.012) but no Timepoint x Group interaction (F(6,225) = 0.905, p = 0.492). We also did not find any group effect (F(2,75) = 1.561, p = 0.217). Similar to MT2, variance was similar across groups at MT3.

To account for any noise created by participants having different variabilities at baseline, we subtracted the participants’ Pre value from each of their Post values. A mixed model ANOVA with three groups x three timepoints on this baseline-adjusted data showed that at MT1, the most challenging speed, there was a significant main effect of Timepoint (F(2,150) = 12.575, p<0.001) but no interaction effect of Timepoint x Group (F(4,150) = 0.838, p = 0.503). There was a significant group difference (F(2,75) = 3.560, p = 0.033), suggesting that after adjusting for individual differences at baseline, movement variability at MT1 improved differently for different groups. Tukey’s post-hoc test revealed that the Sham group improved movement variability significantly more than the DLPFC group (p = 0.027) at MT1. At MT2, which was the training speed, we found a main effect of timepoints (F(2,152) = 10.659, p<0.001) but no Timepoint x Group interaction (F(4,152) = 1.067, p = 0.375) or group effect (F(2,76) = 0.687, p = 0.506). Variability improved similarly across groups at the training speed. At MT3, which was the slowest speed range, the timepoint main effect was significant (F(2,150) = 5.992, p = 0.003) but there was no interaction effect of Timepoint x Group (F(4,150) = 1.203, p = 0.312). We also did not find any significant group effect (F(2,75) = 0.423, p = 0.657) at MT3, meaning at the slowest speed all the groups improved similarly in variability across timepoints.

#### Movement Accuracy

The mixed model ANOVA with three groups x four timepoints on movement accuracy suggests that at speed range MT1, there was a significant main effect of timepoints (F(3,228) = 6.710, p < 0.001) but no Timepoint x Group interaction (F(6,228) = 0.971, p = 0.446). There was a group effect (F(2,76) = 4.691, p = 0.012), with DLPFC groups accuracy being significantly higher than SMG (p = 0.008). In other words, the groups generally improved across timepoints, with DLPFC group being consistently better across all timepoints on speed range MT1. At speed range MT2, we found a significant main effect of timepoints (F(3,228) = 11.004, p < 0.001) but no Timepoint x Group interaction (F(6,228) = 0.884, p = 0.507). There was a borderline group effect (F(2,76) = 3.019, p = 0.055), with DLPFC group’s accuracy being significantly higher than SMG (p = 0.043). On MT2 speed range, learning improved across all timepoints for all the groups. At speed range MT3, we also found a significant main effect of timepoints (F(3,228) = 12.366, p < 0.001) but no Timepoint x Group interaction (F(6,228) = 0.616, p = 0.717). We did not find any between subject group effect (F(2,76) = 2,095, p = 0.130).

To adjust and correct for individual differences in baseline (Pre timepoint) performance, we subtracted accuracy at the Pre timepoint from each Post timepoint and performed a mixed-model ANOVA on the baseline-adjusted performance with three Groups X three Timepoints (Pos1, Post2, Post3). After adjusting for baseline differences, at MT1 we found a main effect of timepoints (F(2,152) = 6.096, p = 0.003) but no Timepoints x Group interaction (F(4,152) = 0.307, p = 0.873). There were no group differences (F(2,76) = 2.270, p = 0.110) at MT1 meaning the groups improved across timepoints but there were no differences in performance between the groups even though Sham seemed to perform better. For speed range MT2, there was a timepoint main effect (F(2,152) = 10.812 p<0.001) but no Timepoint x Group interaction (F(4,152) = 0.167 p = 0.995). There was also no between-subject group effect at speed range MT2 (F(2,76) = 2.093, p = 0.130). Similar to MT1, there were no group differences found at MT2 speed range, but the groups’ performance improved across the timepoints. MT3 also had a main effect of timepoint (F(2,152) = 14.888, p<0.001) but no interaction effect of Timepoint x Group (F(4,152) = 0.258, p = 0.904). There were no between-subject group differences either (F(2,76) = 1.173, p = 0.315), meaning accuracy across groups improved similarly even after correcting for baseline differences.

### Proprioception

#### PSE

We performed a mixed-model ANOVA with three groups (DLPFC, SMG, SHAM) x two timepoints (Pre, Post1) on proprioceptive bias (PSE). There was no significant Timepoint X Group interaction (F(2,76) = 1.153, p = 0.321) or main effect of timepoint (F(1,76) = 1.525, p = 0.221) or group (F(2,76) = 1.728, p = 0.185), indicating that bias was similar across timepoints and between groups (Fig. 6A). DLPFC group’s performance seemed to worsen from Pre to Post but was not significantly different from baseline (p= 0.370) either. Individual performance variability was high (Fig. 6B).

**Figure 4.**
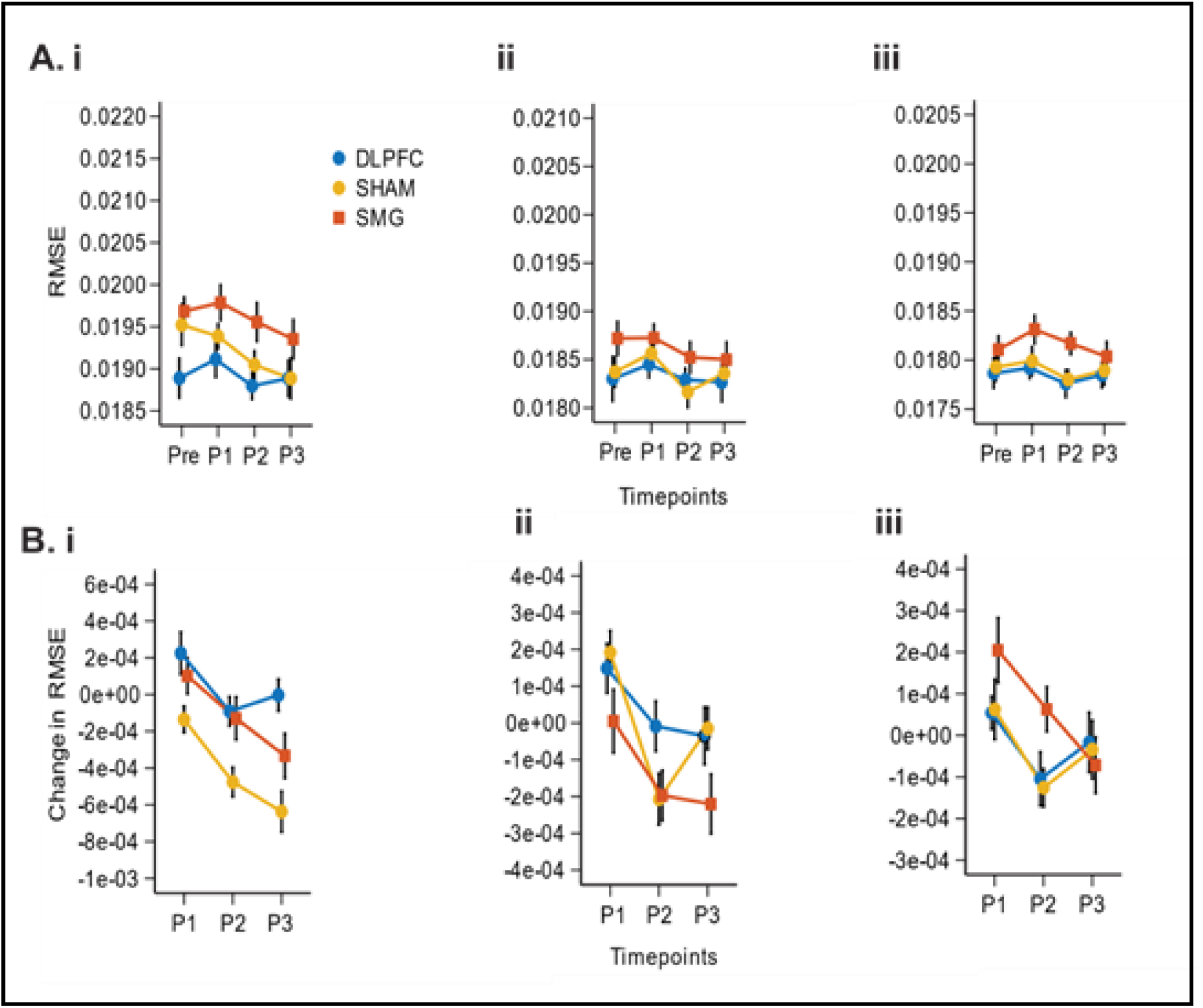
Changes in movement variability (RMSE) from Pre to post-intervention timepoints (P1-P3) at each speed range (MT1, MT2, MT3). Lower values reflect better performance. **A. i.** RMSE during MT1 speed range for the three groups. DLPFC group started off with less variability than the control and SMG group at baseline. **ii.** & **iii** Movement variability (RMSE) during MT2 and MT3 speed ranges respectively for the three groups. **B.** Baseline adjusted performance (All Post timepoints – Pre) at each speed range. **i.** For MT1, variance decreased from Post1 to Post3 for the Sham group as compared to the intervention groups. **ii** & **iii** Baseline adjusted variance at MT2 and MT3 speed ranges. There were no significant group differences.

**Figure 5.**
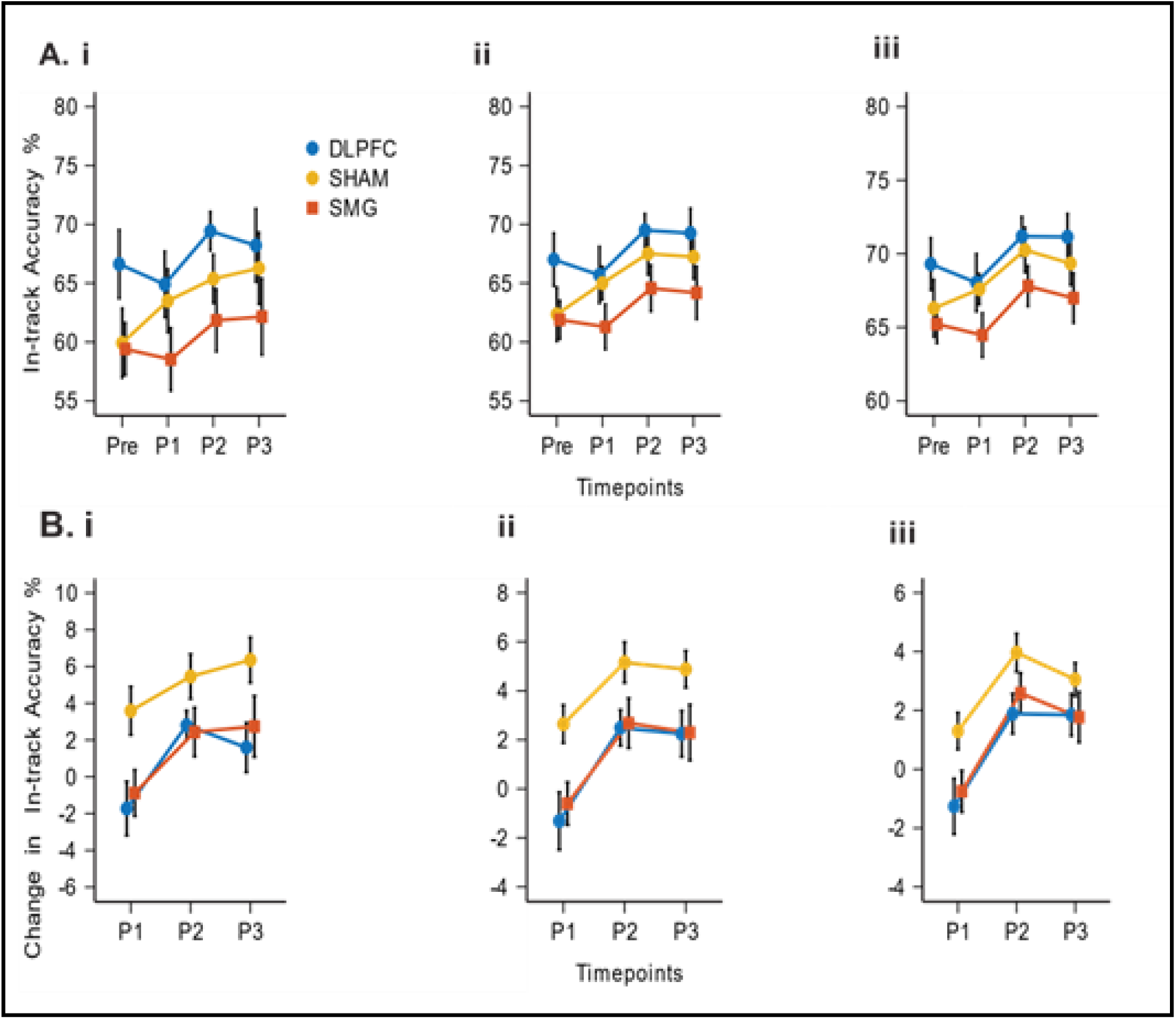
In-track accuracy changes from Pre to Post3 timepoints at each speed range. **A. i.** In-track accuracy at MT1. DLPFC group’s performance was significantly higher compared to the SMG group. **ii.** In-track accuracy at MT2. There was a borderline group effect with DLPFC group’s performance significantly higher compared to the SMG group. **iii.** In-track accuracy at MT3. **B.** Baseline adjusted accuracy performance (All timepoints – Pre) at each speed range (i-iii). There were no significant group differences.

**Figure 6.**
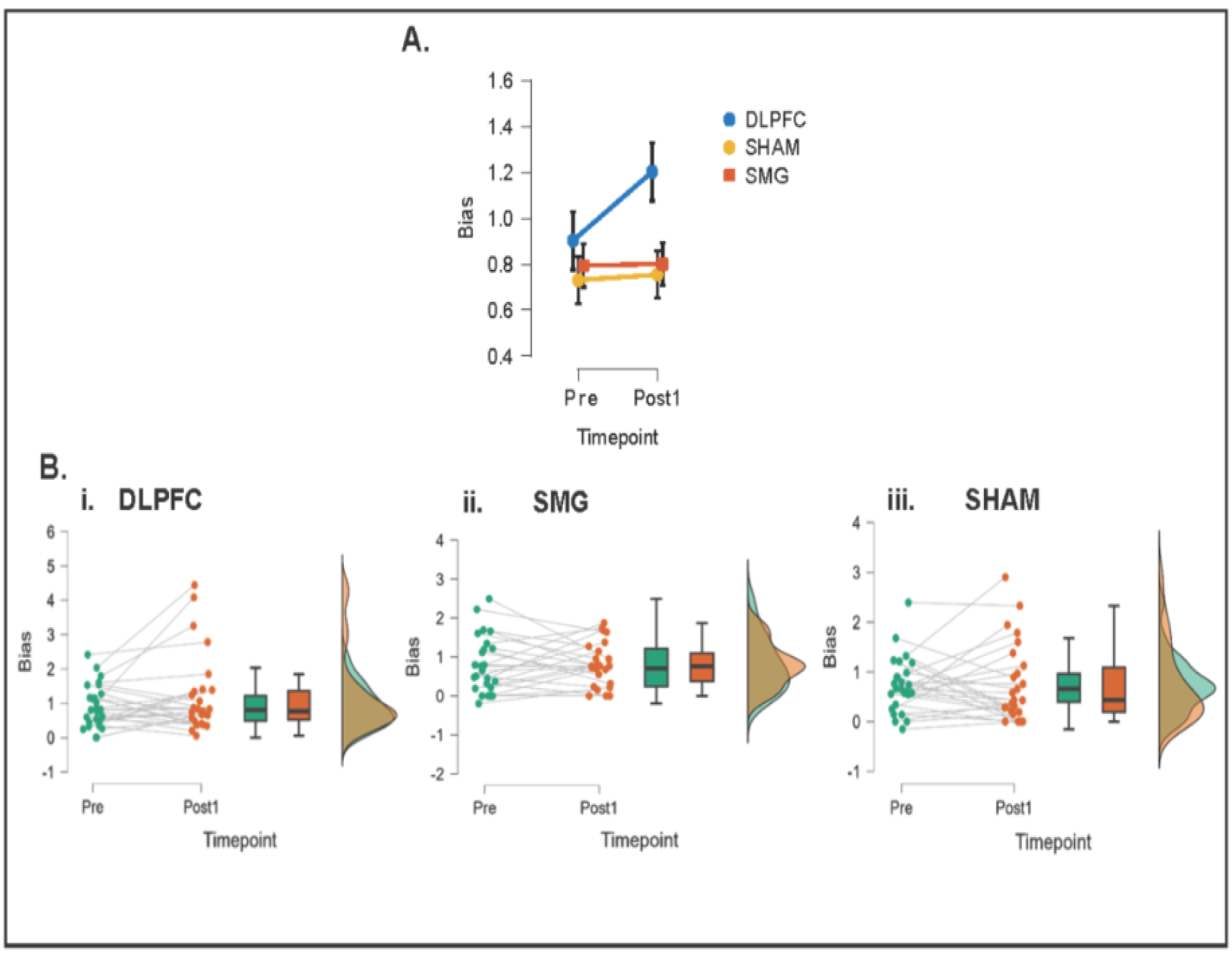
Proprioceptive bias changes from Pre to Post intervention. Lower values represent better proprioceptive performance. **A.** Mean proprioceptive bias magnitude seemed to increase (worsen) more from Pre to Post1 during the DLPFC group as compared to the SMG or Sham (Control) group but was not statistically significant. There were no group differences or interaction effects in proprioceptive bias. Error bars represent the standard errors of the mean. **B. i**. Each line represents individual subjects’ performance from baseline to Post1 in the DLPFC group. **ii.** Each line represents individual subjects’ performance from baseline to Post1 in the SMG group. **iii.** Each line represents individual subjects’ performance from baseline to Post1 in the SHAM group.

#### JND

A mixed-model ANOVA was performed on proprioceptive sensitivity (JND), with three groups x two timepoints. There was no significant Group X Timepoint interaction (F(2,75) = 0.322, p = 0. 726) or main effect of Timepoint (F(1,75) = 3.319, p = 0.072) or group (F(2,75) = 0.135, p = 0.874), indicating that sensitivity was similar across timepoints and between groups (Fig 7A). Every group’s performance seemed to worsen from Pre to Post but no significant difference from baseline was found. Individual performance variability was high (Fig 7B).

**Figure 7.**
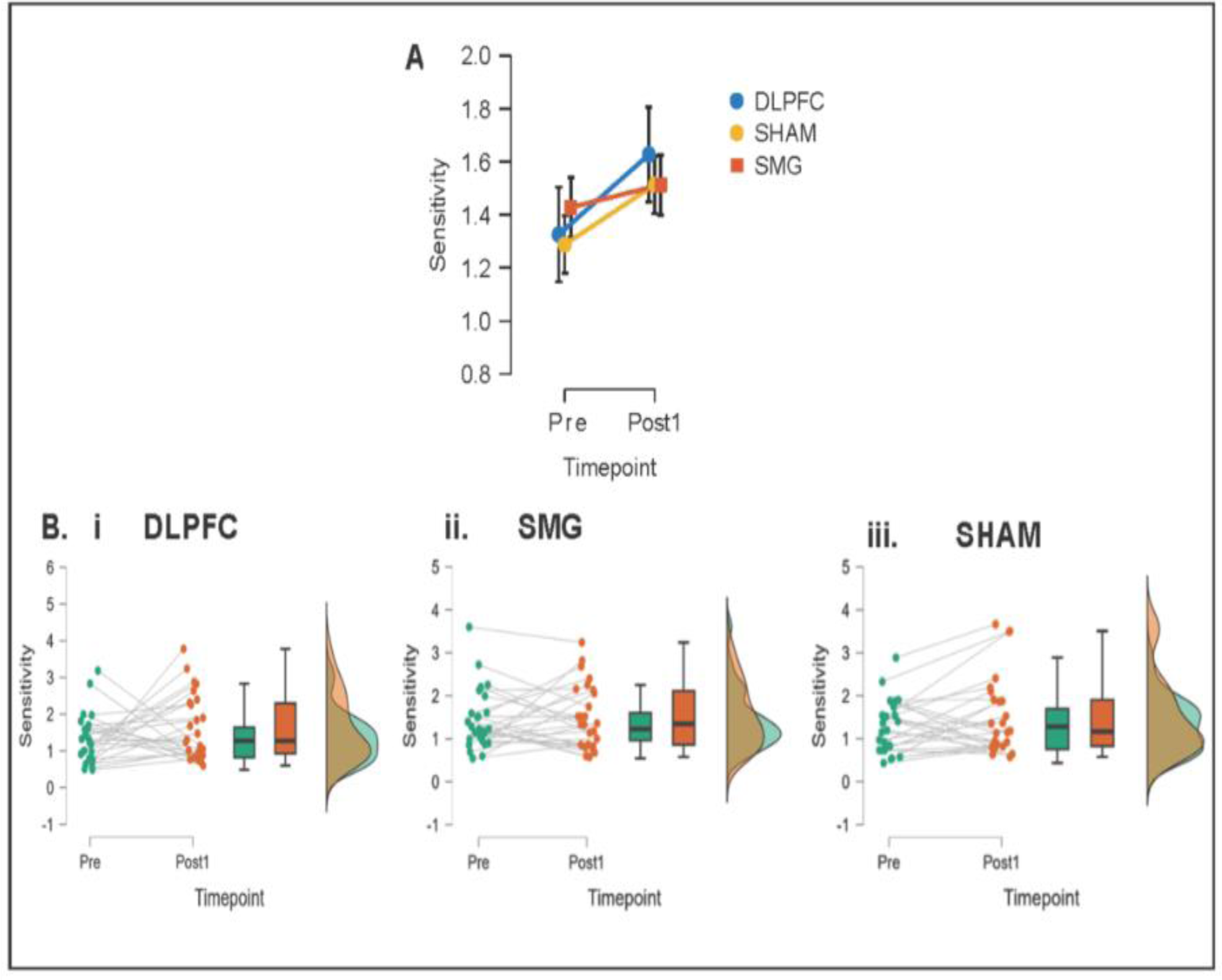
Proprioceptive sensitivity changes from baseline to post intervention (cTBS). Lower values represent better proprioceptive performance. **A.** Mean proprioceptive sensitivity magnitude from Pre to Post1. There were no group differences or interaction effects in Proprioceptive sensitivity. Error bars represent the standard errors of the mean. **B. i-iii**. Each line represents individual subjects’ performance from baseline to Post1.

### Subjective ratings

Subjective ratings of attention, fatigue, sleep and pain were similar across groups. There was no significant difference between the baseline RMT either (Table 1). There was a main effect of group for the handedness scores (F(2,76) = 3.193, p = 0.047). Post-hoc Tukey’s test revealed that the DLPFC group was significantly different from the Sham group (p = 0.038) meaning that the DLPFC group participant were more strongly right-handed as compared to the control group.

**Table 1.** Summary of neurophysiology and subjective ratings across groups.

| | **DLPFC**<br>(mean ± 95% CI) | **SMG**<br>(mean ± 95% CI) | **Sham**<br>(mean ± 95% CI) |
| --- | --- | --- | --- |
| RMT (% MSO) | 49.33 ± 3.8% | 50.73 ± 3.5% | 50.31 ± 3.2% |
| Attention rating | 8.04 ± 0.49 | 7.69 ± 0.48 | 8.04 ± 0.45 |
| Fatigue rating | 3.82 ± 0.68 | 3.92 ± 0.67 | 3.58 ± 0.56 |
| Sleep rating | 7.44 ± 0.56 | 7.69 ± 0.55 | 7.85 ± 0.58 |
| Pain rating | 1.74 ± 0.48 | 1.73 ± 0.39 | 1.27 ± 0.24 |
| Handedness | 89.18 ± 4.22 | 85.22 ± 6.66 | 78.39 ± 7.64 |

RMT: Resting motor threshold. MSO: Maximum stimulator output. All ratings were on a scale from 1-10, reported verbally, with 10 being the best attention and sleep and the worst fatigue and pain.

## DISCUSSION

In this study, we investigated the contributions of two different brain regions, the dorsolateral prefrontal cortex (DLPFC) and the supramarginal gyrus (SMG), during motor skill learning and proprioception, and compared these findings with an inactive sham group. We also assessed whether motor training, both short- and long-term, would improve performance. We found that all three groups demonstrated improvements in movement accuracy and variability indicating that they successfully learned the maze tracing skill. However, movement variability in the DLPFC and SMG groups improved less compared to the sham group during the speed-accuracy tradeoff at the fastest speed range. This suggests that the DLPFC and SMG may play a role in the more challenging aspects of skill learning. This finding aligns with the established associations of these regions with higher-level cognitive functions (Goto et al., 2011) and proprioceptive processes (Ben-Shabat et al., 2015).

### DLPFC and SMG involved during more complex motor skill

Motor learning can take a variety of forms, such as learning to improve reaction time, learning finger tapping patterns, and adjusting movements to compensate for perturbations, which is known as motor adaptation (Bastian, 2008; Krakauer et al., 1999, 2000). In contrast, motor skill learning requires improving a movement pattern without any perturbation (Shmuelof et al., 2012). Time taken to improve or reduction of reaction time on a skill is one way to define complexity, but another crucial aspect is the kinematic complexity of the movement pattern involved. Much of the literature in this area has examined either motor adaptation or simple motor skills, e.g., finger button presses or planar straight reaches (Dayan & Cohen, 2011; Ostry & Gribble, 2016; Wong et al., 2011). Less is known about learning movement patterns that are more demanding kinematically. For example, a movement pattern with abrupt turns would require a series of coordinated movements by different joints in order to accurately perform the pattern, such as the maze task used previously in our lab (Mirdamadi & Block, 2020) and in the present study.

An imaging study by Kami et al. (1995) reported that when participants repeatedly practiced a sequence within a short period of time, it caused an activation in a smaller M1 area initially (‘habituation’) which was followed by an increase in the area of activation (‘enhancement’) (Kami et al., 1995). Kelly and Garavan (2005) connected this shift in activation to the reduced dependence on executive functions and attentional resources that takes place as learning progresses (Kelly & Garavan, 2005). They suggest that fast motor skill learning is mainly mediated by the cortical networks while the long-term learning is mediated by the cortico-subcortical network activation which are known to be involved in “automatic” movements (Floyer-Lea & Matthews, 2004). Learning motor skills necessitates the interaction of several high-level cognitive processes with low-level sensorimotor mechanisms (Krakauer & Mazzoni, 2011). Early learning stages are known to activate the frontoparietal cortices like dorsolateral prefrontal cortex, primary motor cortex, and posterior parietal cortex, while later stage relies more on the subcortical structures like the cerebellum and basal ganglia (Floyer-Lea & Matthews, 2005; Halsband & Lange, 2006). The two regions examined in the present study, dorsolateral prefrontal cortex (DLPFC), part of the frontal cortex and supramarginal gyrus (SMG), a part of the posterior parietal cortex are thus expected to be most important in the early stages of motor learning. The maze tracing skill was practiced in two intervals, the first shorter than the second; however, given that both were on the order of minutes, the experiment likely examined only the early stages of motor learning.

The left DLFPC is known to contribute to working memory/manipulation during skill learning (sequence finger-tapping task) and cTBS over the left DLPFC inhibited improvement in motor skill learning during high working memory demand conditions requiring more dependence on attentional resources and executive function (Lin et al., 2022).The maze task used in this study, is also a complex skill with irregular shape, 1.5 cm wide and performed on different speed ranges, which could make it more working memory/ cognitively demanding, especially at the fastest speed range. It is possible that executing the fastest movements required the brain to constantly adjust the movements to maintain precision at faster speeds making it the most cognitively challenging (Missenard & Fernandez, 2011). Our results were consistent with the previous finding in that movement variability performance failed to improve in the DLPFC group as compared to the Sham group during the skill assessment at the fastest speed range which was 300-600ms. Movement variability has also been studied in the context of running speeds (Wang et al., 2021). Speed is known to significantly influence variability in movement where faster speeds exert more task demands to the system, and more resource allocation is needed to accomplish the task goal.

The SMG group, similarly, did not improve movement variability as much as the Sham group at the fastest speed range. This could be linked to the SMG’s role in proprioception and spatial acuity. In 1982, Ungerleider and Mishkin proposed parietal cortex to be a part of the dorsal visual stream which is involved in spatial perception (Mishkin et al., 1983). More recently, Pollock and colleagues (2020) examined 54 healthy individuals who performed a serial reaction time task (SRTT). They applied anodal, cathodal, and sham tDCS over the left PPC either before, during or after training and found that inhibiting left PPC interfered with the replication of sequences learned previously (Pollok et al., 2020), suggesting that PPC contributes to motor sequence learning. Previous studies have also shown PPC’s engagement in learning new motor skills and its role in transmitting information from the sensory cortex to motor cortex once the learning has been consolidated (Karabanov et al., 2012). In addition, PPC is important for processing higher-level proprioceptive information. Inhibiting PPC is shown to interfere with the reproduction of sequences which have already been acquired and not during online learning (Pollok et al., 2020). Findlater and colleagues’ study on stroke patients revealed the crucial involvement of PPC, especially right SMG, in processing the sense of position (Findlater et al., 2016). Right SMG is a part of the inferior parietal lobe which is associated with spatial processing and multisensory integration (Ben-Shabat et al., 2015). Right SMG is also known to be crucial for integrating proprioceptive information with spatial frameworks which is essential for a person to perceive body position and movement in a spatial reference frame (spatial perception) (Ben-Shabat et al., 2015). The maze skill task in this study had several abrupt turns which might have made it more spatially challenging, especially at the fastest speed making it consistent with the SMG’s involvement in movement variability. We also found no effect of SMG on movement accuracy which was consistent with other brain areas such as primary and secondary motor areas being more crucial for movement accuracy (Durand-Ruel et al., 2023).

### Effect of motor learning on proprioceptive function

Several studies have shown that motor learning is associated with changes in proprioception. In 2009, Cressman & Henriques revealed that visuomotor adaptation changes our sense of position in the hands (Cressman & Henriques, 2009). Learning a directional force-field adaptation causes systematic changes in our perception of hand position in the form of bias in the direction of the load/ force that is learned (Ostry et al., 2010) and learning to use precise hand movements enhances limb position sense (Wong et al., 2011). This led to the hypothesis that the learning process to facilitate new behavior includes both motor and sensory changes (Haith et al., 2008; Vahdat et al., 2011). Another study by Wong et al, 2012 had subjects perform a task recreating a particular hand path, either a word which was handwritten or a circle at constant speed. The required movements were demonstrated visually to the subjects regularly through the training. A group of subjects were given additional proprioceptive information of the trajectories to be performed. This was done by a robot passively moving their hand through the desired trajectories with visual information given about the hand’s desired position. They found that the group of subjects that were provided with additional proprioceptive training learned the desired movements better (Wong et al., 2012). A review showed converging evidence that proprioceptive training overall results in 52% enhancement of motor function throughout every outcome measure (Aman et al., 2015). For example, 30 Hz or higher muscle vibrations for long durations mediated up to 60% improvements, and training on reaching targets and joint position improved 48% of the position sense in joints (Aman et al., 2015). These results suggest that proprioceptive training stimulates cortical reorganization, which strengthens the notion that proprioceptive training can be used in neurorehabilitation to improve motor functioning.

It had been shown that motor learning or training improves not only trained motor skill but also proprioceptive function (Ostry et al., 2010). Some studies have employed active movement training and exercise with visual feedback and found substantial improvements in proprioception (Beets et al., 2012; Elangovan et al., 2017). In contrast, there have been studies that found gains in motor function but not in proprioception such as in patients with ACL reconstruction after neuromuscular training. They found that training helped improve knee function after ACL reconstruction surgery but there was no improvement in proprioception after neuromuscular training (Risberg et al., 2007). Our results were consistent with the latter studies where we found that motor training helped in improving the movement variability or motor performance across timepoints but had no effect on proprioception. This suggests that the effectiveness of motor training might differ based on the type of training performed. This could be critical for rehabilitation purposes where the same kind of training might not work for all patients; individualized training may be better due to high variability in proprioceptive performance.

A previous cTBS study from our lab found the control group’s proprioceptive sensitivity enhanced after training and retention (Mirdamadi & Block, 2021). They applied cTBS over primary somatosensory cortex (S1) and cerebellum (CB) and found impaired proprioceptive sensitivity only in the horizontal dimension and not sagittal. Differences were found during different phases of learning, where S1 contributed to online proprioceptive decrement that persisted at retention and CB contributed to offline proprioceptive decrements (Mirdamadi & Block, 2021). Whereas, in another study they found that the proprioceptive sensitivity improved in the sagittal dimension after motor practice (Mirdamadi & Block, 2020). The mixed results might indicate that the timeframe and dimension–either horizontal or sagittal–in which the task was performed could show different changes in proprioceptive sensitivity. The results from the present study used a similar maze task for assessments but there were some key differences for testing the hand position sense. They used a passive two-alternative forced choice task, where the participant had to report where their hand was in relation to a visual reference marker; this means a high-level multisensory representation (comparing visual location with proprioceptive location) was assessed (Wali & Block, 2024). In other words, this task would lead to integrating and weighing multiple sensory modalities such as proprioception and vision together. Whereas in the current study the proprioceptive task did not have a visual reference marker, making it a purely proprioceptive assessment of body representation as only one modality was concerned (Wali & Block, 2024). This type of assessment may be a noisier measure, perhaps due to higher concentration requirements. Future studies should investigate how these high-level multisensory representative assessments differ from the mid-level purely proprioceptive assessments and measure the changes in proprioceptive functions over time.

### DLPFC and SMG showed no involvement in proprioceptive function

Increased activity in the sensorimotor cortex, possibly through the fronto-striato-thalamic pathways, after applying anodal tDCS over DLPFC, suggested that DLPFC might be involved in enhancing proprioception in healthy individuals (Beck et al., 2019). It was a between subject study, where each participant completed the stimulation and control conditions both one week apart. The results indicated a decrease in variability in spatial error (lower distance travelled variability) in the group that was given anodal tDCS and performed the task where they had to move the cursor on the monitor using a mouse from the starting position to the center of one of the three target squares with their non-dominant hand either with or without visual information. During the without visual feedback condition, the participants had to solely rely on their proprioception. The sham group did not show a decrease in variability either with or without vision. They concluded that the anodal tDCS over left DLPFC could be used to modulate sensorimotor excitability and thus influence proprioception in the task that require precise movement control without vision. In our study, we modulated the different brain regions by inhibiting them using cTBS and found that the results of proprioceptive function were not in consensus with the previous study. We did not find any left DLPFC involvement in proprioception. One of the reasons this could be is the use of different modulation techniques in the two studies. TMS is usually more focal, and targets more specific brain regions as compared to tDCS which alters a larger brain area because of the spread of the current under the electrodes (Polanía et al., 2018). This means that there are higher chances of anodal tDCS exciting areas around DLPFC.

Studies by Findlater and colleagues used a passive position sense task (arm position matching) and statistical lesion-behavior analysis to determine which brain lesions were most associated with deficits in movement and proprioception in patients with stroke. Their findings identified posterior parietal cortex; specifically, supramarginal gyrus (SMG) as a critical brain region in the neural network associated with our position sense. They found the SMG damage was associated with proprioceptive deficits in some patients, suggesting the region’s contribution to perception and position sense (Findlater et al., 2016, 2018). An fMRI study by Ben-Shabat and colleagues found that right SMG had significant activation during proprioceptive assessments in heathy people. They also tested some stroke patients with proprioceptive impairments and observed decreased activation in the right SMG (Ben-Shabat et al., 2015). On the other hand, our findings suggest that right SMG inhibition did not disrupt proprioceptive functioning (Bias or sensitivity). This could be due to the difference in the type of proprioceptive assessments used since SMG is known to integrate multiple sensory modalities which helps us in limb position sense and spatial awareness making it to be involved in high-order proprioceptive processing (Findlater et al., 2016) whereas as mentioned earlier our task was purely proprioceptive (Wali & Block, 2024). This suggests that the level of proprioceptive function might have different neural network distribution involved. Future studies could assess this difference in high-order, mid and low-level proprioceptive functions, and their associated neural networks.

### Limitations and future directions

Fatigue is known to impair task performance and motor skill learning (Branscheidt et al., 2019). Each participant took approximately two hours to complete the study, which could have resulted in fatigue and poor attention during or by the end of it, even with breaks. This likely contributed to the substantial variability observed in individual task performance, especially during proprioceptive assessment which required subjects to be very focused. Due to random assessment ordering, some participants performed the skill assessment task first and others performed the proprioceptive assessment first. We already know that the effect of cTBS lasts for up to 60 minutes but there is no consensus as to when the strongest effect is observed (Jannati et al., 2019; Wischnewski & Schutter, 2015) and at which assessment timepoint. This was likely another source of variability in the results. Finally, we also found that the handedness scores between the DLPFC group and the control group significantly differed, which could have resulted in the baseline differences during motor skill accuracy and movement variability assessment.

Future research is needed to test further the involvement of Dorsolateral Prefrontal Cortex and Supramarginal Gyrus and their contributions to motor skill learning and proprioception. Given that DLPFC and SMG showed less improvement in movement variability on a speed-accuracy tradeoff assessment task at fastest speed, it would be intriguing to investigate the contribution of these brain regions in different task complexities with more complex (faster speeds or narrower maze) or simpler motor task (slower speeds and wider maze), examining the effects during different stages of learning such as consolidation and retention. Further studies can also compare variations in workspace, such as testing in the horizontal plane rather than sagittal, and investigate the differences between low, mid and high-order proprioceptive processing by comparing multiple assessments.

### Conclusions

Results suggest that improving performance on a maze tracing skill at a fast speed range may require resources from both DLPFC and SMG during movement variability but not accuracy. There was no evidence of a role for these regions in proprioceptive changes.

## Author Contributions

MW: Conceptualization, Methodology, Formal Analysis, Investigation, Writing – Original Draft, Writing – Review & Editing, Visualization. HJB: Software, Writing – Original Draft, Writing – Review & Editing, Visualization, Supervision, Funding Acquisition.

